# Adaptive radiation during long-term experimental evolution of the multicellular bacterium, *Streptomyces*

**DOI:** 10.1101/2025.08.20.671327

**Authors:** John T Munnoch, Silja Vahtokari, Leena Kerr, Paul A Hoskisson

## Abstract

Adaptive radiation of a single lineage into novel ecological niches underpins the evolution of biodiversity. To study adaptation in the industrially and ecologically important bacterium *Streptomyces,* a long-term evolution experiment (LTEE) was undertaken in a strain lacking several antibiotic biosynthetic gene clusters, facilitating the study of epistasis in antibiotic biosynthesis. *Streptomyces coelicolor* is a filamentous soil organism that does not undergo sporulation under the LTEE conditions, affording the opportunity to study adaptation under relaxed selection on sporulation. Over 3000 generations, replicate populations showed parallel loss of sporulation, morphological transitions to fragmenting hyphae and diversification in carbon source utilisation. Genomic analyses revealed mutations in developmental and metabolic genes, while reintroduction of the actinorhodin cluster demonstrated pervasive negative epistasis affecting antibiotic production. These findings reveal adaptive diversification and antagonistic interactions in regulatory pathways under relaxed selection, providing insights into the evolution of complex multicellular bacteria and informing industrial antibiotic production strategies.

## Background

Adaptive radiation plays a central role in the emergence of biological diversity [1]. It is thought that ecological opportunity is an important means for generating the phenotypic innovations that allow adaptation into available ecological niches, but the underlying mechanisms that drive this are poorly understood [2–4]. Experimental evolution studies have revolutionised our understanding of the fundamental ecology and evolution of adaptation [1,5–9], and the eco-evolutionary dynamics of replicate populations can be highly informative regarding the wider process of adaptation [5,6]. Genetic adaptation to a single environment often results in rapid fitness gains but may result in the loss of fitness in other environments, resulting in ecological specialisation [10]. This adaptative process often leads to a decline in certain traits that are unimportant in the new environment. In this situation, two evolutionary processes appear to predominate: the accumulation of mutations, resulting in genetic drift of those genes not maintained by selection, and antagonistic pleiotropy, such that mutations in one environment may be deleterious in another [10–13]. The underlying evolutionary mechanisms are difficult to distinguish but are likely to be the same regardless of whether they are occurring in the natural environment or via prolonged laboratory culture [14,15]. Moreover, in the natural environment it is increasingly demonstrated that there is interdependency of mutations (epistasis) and this interdependency may act in a synergistic or antagonistic manner [16,17]. This can in part explain the commonly observed phenomenon of a decline in the rate of fitness in certain traits which can increase over time in natural environments and in microbial evolution experiments [10,17].

The filamentous soil bacteria of the genus *Streptomyces* are prolific producers of a range of bioactive natural products and possess a complex developmental lifecycle [18–20]. The lifecycle of *Streptomyces* begins with the germination of a spore which gives rise to an apically growing hypha, which elongate and branch during growth. In response to nutrient limitation or stress, the developmental process of *Streptomyces* is initiated, where aerial hyphae rise into the air, curl and mature into unigenomic spores through the laying down of septa in the aerial hyphae [18]. Growth in shaken liquid culture prevents sporulation in many species of *Streptomyces*, and their normal growth habit is to form aggregates of hyphae that propagate through apical growth and fragmentation [21,22]. Another remarkable feature of *Streptomyces* biology is that they have large (∼8-10 Mbp) linear genomes, with extraordinary catabolic capability [23,24]. *Streptomyces* also possess telomere-like sequences at each end of their linear chromosomes [25] and it is believed that this feature leads to the genetic instability observed in these organisms and results in phenotypic diversity [26–31]. Attempts to explain this genetic instability in terms of the ecology and evolution of *Streptomyces* has recently demonstrated that it can, at least in part, be explained through division of labour in colonies, particularly in structured, terrestrial environments [20,32–34].

To further understand adaptation in *Streptomyces,* a long-term evolution experiment (LTEE) was undertaken using the model species *Streptomyces coelicolor* to investigate the role of adaptation to a single environment. Over 3000 generations, replicate populations exhibit extensive parallel loss of sporulation (antagonistic pleiotropy) and negative epistasis in antibiotic production. Major morphological transitions also occurred with parallel fragmentation phenotypes emerging within the populations, suggesting adaption to a homogeneous liquid environment along with widespread catabolic adaptation. Sequencing of isolate genomes provides insight into the underlaying genetic causes for the observed phenotypes. This *Streptomyces* LTEE (*S*LTEE) provides a unique system to investigate the role of morphology and development under conditions that are non-permissive for sporulation, enabling the study of antibiotic production independent of development and in an environment analogous to that used in industry to produce antibiotics.

## Methods

### Growth and Maintenance of Bacterial Strains

The progenitor strain used in this study was *Streptomyces coelicolor* M1152 (Δ*act,* Δ*red,* Δ*cpk,* Δ*cda, rpoB* [C1298T]) [35]. All independent lineages were derived from a single colony clone from a spore stock of this strain.

### Growth assays

Each replicate population of *S. coelicolor* M1152 was grown in 250 ml Erlenmeyer flasks containing 50 ml of YEME medium [36]. Cultures were propagated every 72 hours by transferring 0.5 ml of culture into 49.5 ml of fresh YEME medium, using a serological pipette to avoid size selection of growth aggregates. In the 72-hour growth period, cultures attained stationary phase. Growth of populations and isolates was measured using a TECAN Spark plate reader with populations and isolates first grown in appropriate media overnight (YEME or MM medium). Samples were then washed and normalised before being placed in the plate reader in 24 well plates for 24h. Growth rate was then calculated using the formula μ = (ln OD₂ - ln OD₁) / (t₂ - t₁).

### Diversity measurement

Individual clonal samples were obtained for phenotypic examination through plating on to mannitol soya flour agar and individual developmental phenotypes were scored relative to the progenitor strain. Mixed population samples were obtained periodically for future study through freezing each passage of cells (10 ml of culture with 5 ml of 50% glycerol) and these are stored at -80 °C. The Diversity Index (*H*) was calculated from analysis of 100 colonies plated from each independent population at the indicated generation point using the Shannon-Weaver index equation [*H =* (*N* log*N* - Σ*n* log*n_i_*)/*N*].

### Microscopy and cell aggregate size determination

The area of cell aggregates was determined by microscopy, where samples were visualised using a TE-2000 inverted widefield epi-fluorescence microscope with a x20/0.50 numerical aperture objective lens (Nikon, Japan). Phase contrast images were acquired using an ORCA-100 CCD camera (Hammamatsu, Japan) and illumination was provided by a tungsten-halide lamp. Images were acquired using NIS Elements software and data was processed for visualisation using FIJI[37] . Images were scaled (2360 pixels/mm) using a 1 mm graticule (Analyze/Set Scale). Areas were measured by manually drawing round pellets (using freehand drawing tool), measured (Analyze/Measure) and these images flattened and saved (Image/Overlay/Flatten). R scripts were used to visualise the data.

### Metabolic capability

Catabolic capability of the *S*LTEE populations was determined using OmniLog analysis with Biolog plates PM1 (Biolog, 12111). Strains were grown from stocks in liquid culture for 24 hrs at 30°C. Biomass was collected by centrifugation and added to inoculation fluid IF-0a (Biolog 72268) to an O.D_600_ of 0.04, to create the initial cell suspension. Experimental inoculation fluid (per 24 mL) was prepared using 20 mL stock of IF-0a solution, with the addition of the metal ion cocktail (1.2 mL; containing 5 mM each: ZnCl_2_ 7H_2_O, FeCl_2_ 6H_2_O MnCl_2_ 4H_2_O, CaCl_2_ 2H_2_O, filter sterilised). Dye Mix D (0.24 mL; Biolog, 74224) and 1.2 mL of sterile distilled water were added to 2.32 mL of cell suspension to give a final volume of 24 mL. This was inoculated into PM1 plates and incubated at 30°C for 2 days in the OmniLog instrument. Data were extracted using the Biolog software (conversion of D5E to OKA: D5E_OKA Data File Converter v1.1.1.15 and extraction of raw kinetic data using PM analysis software: Kinetic V1.3). Data was processed using BactEXTRACT [38], visualised and compared using R scripts (https://github.com/PaulHoskisson/StreptomycesLTEE/tree/main/5_Biolog_populations/). In brief (all files available on github), Biolog data was input into template Tecan data file for 48 hours of growth (171000 seconds). This was uploaded and results generated for the whole 48 hours. All ‘growthParameters’ data was downloaded. Growth curves were visualised using R (https://github.com/PaulHoskisson/StreptomycesLTEE/tree/main/5_Biolog_populations/4_BactExtract) and those exhibiting classical bacterial growth were used (https://github.com/PaulHoskisson/StreptomycesLTEE/tree/main/5_Biolog_populations/3_plots). Supplementary data available here https://doi.org/10.6084/m9.figshare.29950388.v1

### Actinorhodin assay

The actinorhodin biosynthetic gene cluster (BGC) contained on pAH88 [35] , a BAC derived from SCBAC28G1 was introduced periodically into the population via conjugation from the *E. coli* strain ET12567 (*dam, dcm, hsdS*), containing the driver plasmid pUZ8002 to bypass the methyl-specific restriction system of *S. coelicolor* [36,39]. Actinorhodin concentration was determined according to Feeney et al., [36].

### Genomic DNA extraction

enomic DNA was extracted from *S. coelicolor* using the *Streptomyces* DNA isolation protocol described by Feeney et al. [36].

### Genome sequencing

Short-read genome sequencing of the progenitor strain (*S. coelicolor* M1152) and evolved isolates was carried out using an Illumina NovaSeq 6000 by (Novogene, UK). The *S. coelicolor* M1152 strain was sequenced at the start of the experiment to ensure exact knowledge of the progenitor genotype (Bioproject: PRJNA1304580, with reads available SAMN50561540). The *Streptomyces coelicolor* M1152 genome was constructed *in silico* manually following the publication describing its construction [35] using the *S. coelicolor* M145 reference genome ([23] NCBI: NC_003888.3 [now suppressed by NCBI]) and Snapgene (Version 7.2.1). This reference genome was used as the *Streptomyces* LTEE progenitor reference genome (Bioproject: PRJNA1304580) following SNIPPY analysis (https://github.com/tseemann/snippy) and is available here (https://github.com/PaulHoskisson/StreptomycesLTEE/tree/main/1_Genome_sequencing_M1152_genome). Isolate genomes were analysed using Breseq (version 0.38.3) [40] using the *Streptomyces* LTEE progenitor genome as the reference.

## Results

### The generation of a versatile experimental evolution system for studying adaptive evolution and epistasis in antibiotic production in *Streptomyces*

To investigate how adaptation and epistasis influence antibiotic production, a Long-Term Evolution Experiment (LTEE) with the super host strain *Streptomyces coelicolor* M1152 was initiated (**Fig. 1A**). This strain of *S. coelicolor* used is deficient in production of the blue antibiotic actinorhodin (ACT), the red antibiotic undecylprodigiosin (RED), the calcium dependent antibiotic (CDA), the yellow polyketide antibiotic coelimycin and has an engineered mutation in to *rpoB* to enhance production of antibiotics (Δ*act,* Δ*red,* Δ*cpk,* Δ*cda, rpoB* [C1298T]) [35]. Sequencing and SNIPPY analysis of the progenitor strain *Streptomyces coelicolor* M1152 strain (Bioproject: PRJNA1304580) identified 114 mutations compared to the published *Streptomyces coelicolor* M145 genome [23]. Analysis of the mutations in *Streptomyces coelicolor* M1152 indicate that 38 are intergenic mutations, 17 are non-synonymous, 39 are synonymous mutations and 15 cause frameshifts in coding sequences.

**Figure 1:**
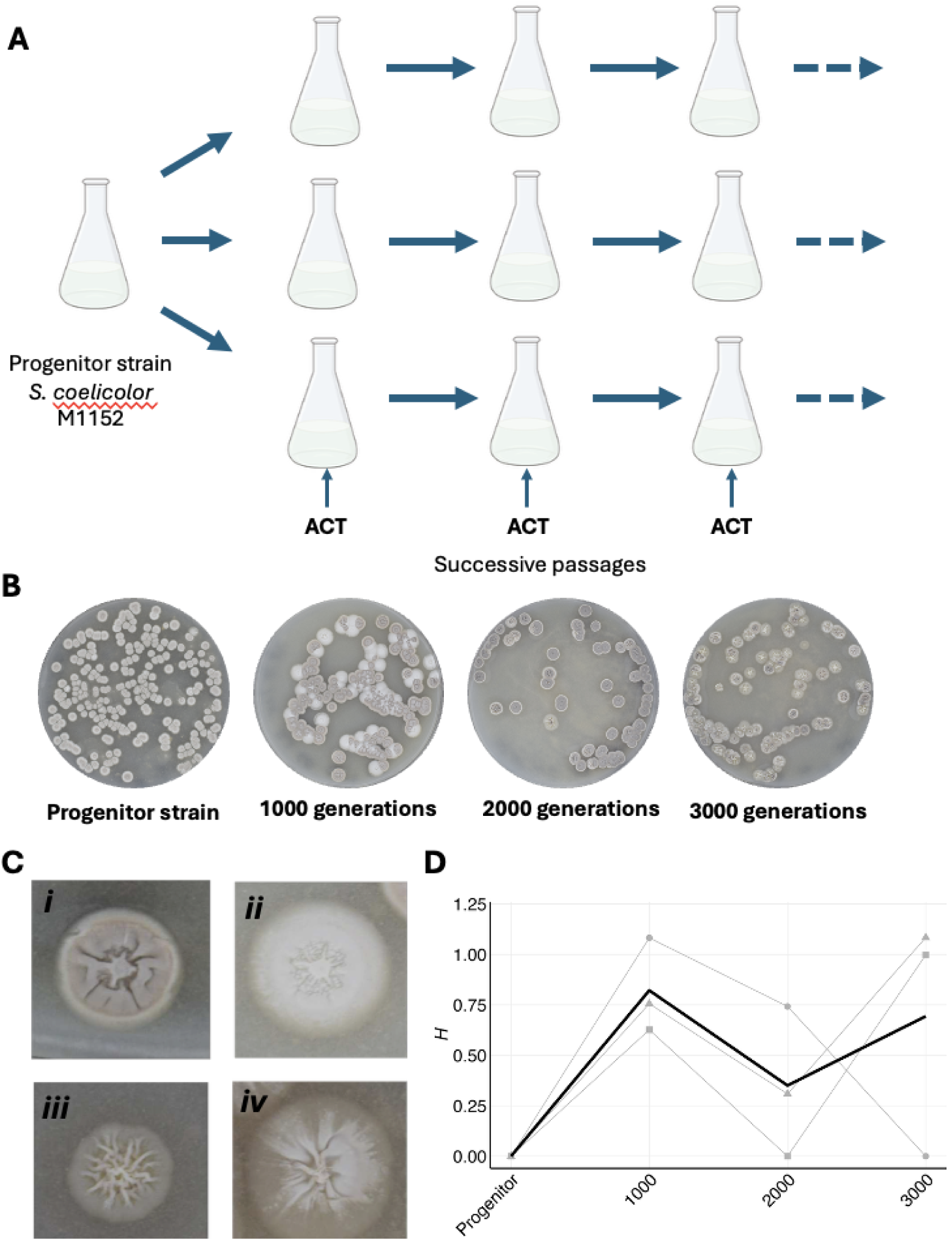
Adaptive radiation of *Streptomyces coelicolor* during long-term experimental evolution (LTEE). **A)**. Experimental set-up of the LTEE showing successive passage of *S. coelicolor* M1152, followed by periodic reintroduction of the actinorhodin (ACT) biosynthetic gene cluster. **B).** Representpulations is indicated by the greative plates demonstrating the phenotypic variation in colony morphotypes in *S. coelicolor* M1152 strains. Replicate populations were founded from a single ancestral grey, sporulating colony and propagated over 3000 generations in YEME medium **C).** Major morphological mutation and transitions occur in the LTEE over 3000 generations and strains can be assigned to four major phenotype groups. ***i*)** Sporulating colonies, phenotypically identical to the progenitor strain ***ii*)** White-aerial hyphae formed, but no spores ***iii*)** Bald – no aerial hyphae or spores formed ***iv*)** ‘Sectored-spreader’ colonies **D).** Colony morphotype diversity during adaptive radiation from the parental grey sporulating morphotype. Diversity was determined through scoring the frequency of morphotypes and using the Shannon-Weaver index. Diversity *(H*) in the three independent replicate populations is indicated by the grey lines (Y1 ●; Y2▲;Y3■) at each major generational point, and the black line indicates the mean diversity *(H*) of the three populations.

Serial passage of three independent parallel populations of the strain in rich medium (YEME; Y1, Y2, Y3), with each passage taking 72 hours (17 generations in the progenitor strain) results in each culture experiencing nutrient sufficiency and nutrient limitation over the course of each passage. This resulted in the cultures reaching stationary phase during every passage, such that the physiological triggers for antibiotic production are experienced by the culture. The deletion of the four main natural product biosynthetic gene clusters (BGCs) results in relaxed selection on antibiotic production, reducing the demand on the cellular pool of metabolites for production of antibiotics. In liquid culture *S. coelicolor* cannot sporulate [41] resulting in relaxed selection on the sporulation too in this experiment. At specific points in the *S*LTEE the ACT cluster was reintroduced into the evolved strains to study epistasis in antibiotic biosynthesis. *Streptomyces* are normally propagated via the collection of spores from mature plate grown cultures [36]. Following repeated passage of the replicate populations via liquid transfer, cultures were periodically plated on to sporulation permissive medium to monitor sporulation in the populations and the identification of developmental mutants.

### Phenotypic divergence of *Streptomyces coelicolor* during long-term experimental evolution

The growth of asexual replicate populations from a single progenitor genotype results in all subsequent genetic variation in the population being generated by mutation [1,42]. The diversification of colony morphotypes within experimental populations of bacteria makes it easy to detect the emergence of heritable genetic diversity [1,43,44]. Given sporulation in *Streptomyces coelicolor* is not permitted in liquid cultures [41], it was hypothesised that relaxed selection on this trait through liquid-to-liquid passage of replicate populations would lead to the emergence of developmental mutants. Developmental mutants are easy to identify in *Streptomyces* with distinct developmental stages falling into two main classes, the bald (*bld*) mutants which are unable to form aerial hyphae and the white (*whi*) mutants which are unable to form mature spores and the grey spore pigment [18,41].

Morphological mutants arise frequently in the *S*LTEE with similar morphotypes emerging rapidly and in parallel across the replicate lines (**Fig. 1B)**. The morphotypes emerging in the *S*LTEE can be classified into four dominant phenotypic groups *i*) Sporulating colonies, phenotypically identical to the progenitor strain *ii*) White strains - where aerial hyphae are formed, but no spores *iii*) Bald strains that form no aerial hyphae or spores *vi*) and colonies that sectored in appearance **(Fig. 1C-*i-iv*)**. The emergence of classical *bld* and *whi* mutations likely reflect the lack of selection for sporulation in the *S*LTEE and the phenotypes persist within the replicate populations.

To quantify the morphotypes in each replicate lineage, the Diversity (*H*) was calculated using the Shannon-Weaver Index [1] **(Fig 1D)**. The homogeneous, grey, sporulating progenitor strain *S. coelicolor* M1152 diversifies within 1000 generations to a maximum point, prior to a decline in diversity as phenotypes fix through adaption to liquid culture **(Fig 1D)**. In two of the independent lineages, there is a second round of diversification from 1000 to 2000 generations which is not seen in the third replicate population. This diversification, followed by selective sweeps of mutants likely leads to the decline in diversity observed in the replicate populations. Whilst this contrasts with observations in homogeneous environments in previous studies using unicellular bacteria [1], the complex hyphal nature of *Streptomyces* likely influences the emergence of diverse morphotypes in liquid culture.

To understand the genetic basis of the morphotypes that emerged within the replicate lineages, 16 isolates were selected that spanned the phenotypic range observed and the three replicate populations. Their genomes were sequenced using an Illumina NovaSeq 6000 by (Novogene, UK) and the data was mapped to the LTEE reference *S. coelicolor* M1152 genome with Breseq. A range of single nucleotide polymorphisms (SNP) were identified (894 mutations across the 16 strains, with 378 unique alleles detected) along with insertions and deletions within the isolates (**Supp. Table S1**). The sequenced isolates from three replicate populations show evidence of the well characterised chromosomal deletions [32, 33] (at the left end of the chromosome resulting in the loss of ∼330 Kbp (loss of genes SCO0001-SCO0326), with one isolate exhibiting the ∼330 Kbp loss at the left-hand end of the chromosome, coupled with a ∼720 Kbp loss at the right-hand end of the chromosome (**Supp. Table S1**). Across the genomes of the 16 isolates from two of the three replicate populations, the average number of mutations per genomes at 3000 generations was 29.85. One replicate (Y2) population (**Supp. Table S1**) exhibited an average of 168.67 mutations per genomes. Examination of the strains from that lineage were found to have a nonsense mutation (W95*; TGG->TGA) in SCO5388 (*nucS*), which is mismatch-repair gene in Actinobacteria, that functions similarly to *mutS* in most bacteria [45]. Mutations in *nucS* have previously been associated with colony sectoring phenotypes [46] similar to those observed in **Fig. 1C*iv***.

Given the developmental and morphological phenotypes that emerge in the *S*LTEE, the sequencing data was interrogated for mutations in the classical *bld* and *whi* genes and other genes known to influence sporulation [18–20] (**Supp. Table S1**). Mutations in *bldH/adpA (SCO2792), bldC (SCO4091), bldKD (SCO5115) ssgF (SCO7153) ssgC* and *(SCO7289)* were present in the 16 sequenced isolates. A total of 11 distinct alleles were identified in the sequenced isolates (in a total of 26 mutations in these genes), indicating the emergence of parallel developmental mutations in the *S*LTEE. Only one isolate showed an absence of mutations in known developmental genes. In the pleiotropic regulator, *adpA* (SCO2792) six independent alleles were identified (one is an intergenic mutation, likely affecting transcription of one or both divergently transcribed genes (lzl13/lzl359; SCO2791 ← / → SCO2792) and was present in all isolates sequenced in one of the replicate populations (Y2; **Supp. Table S1**). Evidence of parallelism is also present with the emergence of two alleles independently in isolates from another replicate population (Y1; L186R and W240*; **Supp. Table S1**). Three further alleles in *adpA* are also found in the Y3 replicate population, with isolates exhibiting either a frameshift (+G at 291/1197 nucleotide or C 6->7 nucleotides at 632/1197 nucleotides) or a nonsense mutation (W97*). The two cell-division associated genes *ssgFC* are found within the large ∼720 kb deletion in the isolate from the Y1 replicate population at 1000 generations.

### Major morphological transitions occur in *Streptomyces* upon serial passage in liquid culture

To understand how *S. coelicolor* adapts to a liquid culture environment, the three replicate populations were examined at 1000, 2000 and 3000 generations by microscopy. The progenitor strain **(Fig 2A)** was found to form the large hyphal aggregates characteristic of *Streptomyces* strains in liquid culture [21,22]. Subsequent serial passage of the replicate populations led to an overall decline in the size of the mycelial aggregates **(Fig 2A & B)**, such that at 3000 generations much of the culture consists of hyphal fragments. This fragmenting phenotype likely represents adaptation to growth in liquid cultures to maximise nutrient acquisition. Hyphal aggregates are known to be nutrient limited resulting in increased stress and cell death at the centre of the aggregates [21,22]. Remarkably, the emergence of this fragmenting phenotype appears to be independent of the developmental phenotypes observed at the same number of generations, where fragmentation phenotype is not always linked to developmental phenotypes (**Fig. 1B, 1C, 1D & Fig. 2A & 2B)**. Examining the genomes of the 16 isolates, all of which were isolated from fragmenting cultures (**Fig. 2A**), possess mutations in the *matAB* locus (SCO2962, SCO2963). Mutation at this locus is known to affect the formation of cell aggregates [21, 22], and is part of a larger locus that appears to be required for cell adhesion and biofilm formation [22]. In isolates from the Y1 replicate population, three independent indels lead to frameshifts in *matAB,* the Y2 replicate population all sequenced isolates contain a SNP that results in a D538H (GAC->CAC) amino acid substitution in MatB, and in the Y3 replicate population all sequenced isolates contain nonsense mutation (Q31*; CAA->TAA) in *matA* (**Supp. Table S1**). The pervasive nature of *matAB* mutations and the fragmenting phenotype strongly suggests a role for these changes in the emergence of the fragmenting phenotypes in the *S*LTEE.

**Figure 2:**
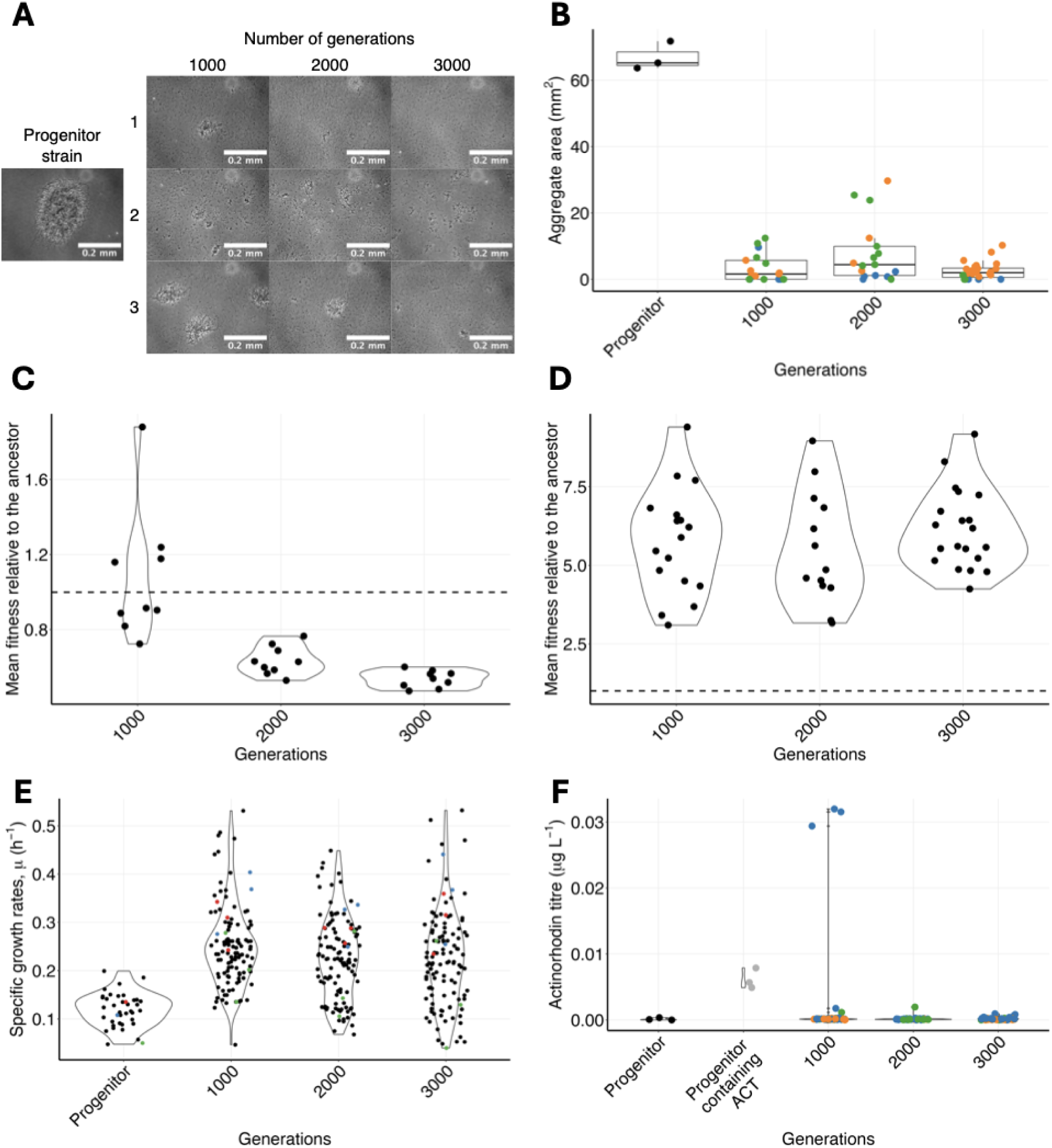
Major metabolic and morphological transitions occur during long-term experimental evolution (LTEE). A). Strains adapt to liquid growth by independently transitioning from an aggregating phenotype in the progenitor to a fragmenting phenotype in all replicate populations. **B).** Quantification of cell aggregate size of *S. coelicolor* M1152 replicate populations shows a decline in mean aggregate area over 3000 generations during adaptation. **C).** Fitness of the replicate populations over 3000 generations. Each point represents the ratio of the growth rate of the evolved population relative to the progenitor strain (*S. coelicolor* M1152) measured in triplicate for each of the three replicate populations. Dashed line represents equivalent fitness. **D).** Fitness of three randomly chosen independent isolates over 3000 generations from the replicate lineages. Each point represents the ratio of the growth rate of the evolved isolate relative to the progenitor strain (*S. coelicolor* M1152) measured in triplicate. Dashed line represents equivalent fitness. **E).** Diversification of carbon source utilisation in the replicate populations over 3000 generations. Each point represents the growth of one of the replicate populations on a particular carbon source in the LTEE. Three carbon sources were chosen to illustrate different usage profiles as the population adapts to the medium through changes to specific growth rate, µ (h^-1^; Further examples in the supplementary data). 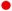 glucose; 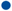 malate; 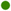 pyruvate **E).** Negative epistasis is pervasive in ACT production in the LTEE. The progenitor strain *S. coelicolor* M1152 makes no actinorhodin, this is restored when the ACT BGC is reintroduced to the strain. Many strains that had the ACT BGC show reduced actinorhodin production relative to the progenitor strain containing ACT. Colours represent independent isolates – three isolates from three independent replicate populations.

### Relative growth rate increases in individual isolates during adaptation in *Streptomyces*

In previous bacterial LTEEs an increase in growth rate has been shown to account for much of the fitness gains observed at the early stages of adaptation [14,47]. To determine if the specific growth rate (µ) increases in the *S*LTEE the populations of *Streptomyces,* as a proxy for fitness. Strains were grown in YEME medium and specific growth rates (µ) were calculated, these were then normalised to the specific growth rate of the ancestor (progenitor) strain to calculate strain and population fitness relative to the ancestor (**Fig. 2C & D;** [48]).

The mean fitness of the populations at 1000, 2000 and 3000 generations was found to decrease as the populations adapt to the liquid culture conditions (**Fig. 2C**). This likely represents increased competition between strains and suggests that adaptive radiation is occurring withing the cultures. To investigate variation within the populations, independent isolates were selected from each replicate population at 1000, 2000 and 3000 generations and the fitness relative to the ancestor determined. Within the isolates, when they are not competing with another strain mean fitness increases steadily relative to the ancestor (**Fig. 2D**). Another notable feature of this growth is there is a general convergence of relative fitness in the strains over 3000 generations.

### Adaptive diversification of carbon catabolism occurs in the *Streptomyces* LTEE

To understand the process that may be responsible for the increase in growth rate in individual isolates, the progenitor and each replicate population was subjected to growth in Biolog phenotypic microarray PM1 plates. The specific growth rate (μ) was calculated for each population in 95 carbon sources and plotted up to 3000 generations (**Fig. 2E)**. Adaptive radiation in the replicate populations results in diversification of the specific growth rate (μ) in different carbon sources over 3000 generations. There is an overall increase in mean growth rate of the replicate populations as they adapt to the medium, for example growth on glucose, malate, proline, and N-acetyl- glucosamine exhibiting higher growth rates than the progenitor strain (**Fig. 2E & Supp. Data 1 & 2**). There are also several examples where replicate populations show decay of catabolic function for a particular carbon source such as pyruvate, tween 40, threonine and glucuronic acid (**Fig. 2E & Supp. Data 1 & 2 and Supp. Fig. 1**).

Whilst it is difficult to pinpoint specific mutations from the sequencing data that may contribute to the increase in relative growth rates and the changes in growth rate on specific nutrient sources in strains there are a number of potential candidates and scenarios, including genome streamlining/loss of catabolic function due to the large chromosomal deletions. In addition, there are a number of mutations in putative transporter genes, solute-binding protein encoding genes, amino-acid biosynthetic genes, cell division genes and metabolic/energy generation genes present in the isolate sequencing data (**Supp. Table S1**). There are also mutations found in regulatory proteins and many mutations in hypothetical protein encoding genes (**Supp. Table S1**).

These data suggest that metabolic diversification is occurring within the populations as strains use different physiological pathways to consume resources. Given that few of the mutations are present in all strains, it suggests that sympatric diversification is occurring within the populations.

### Negative epistasis is pervasive in antibiotic production during long-term experimental evolution

The use of the *S. coelicolor* M1152 superhost strain [33] in the *S*LTEE affords the opportunity to study epistasis in antibiotic production. The replicate populations in the *S*LTEE are subjected to the nutrient limiting conditions that trigger antibiotic production in cultures [49,50], but lack the metabolic burden associated with biosynthesis. Serial passage of *S. coelicolor* M1152, which lacks the ACT BGC enables direct comparison of ACT production in the progenitor strain and the evolved strains following reintroduction of the ACT BGC. Using the isolates from the growth experiments, the ACT BGC [35] was introduced to the strains and relative ACT production assessed in in the cultures. No ACT production was detected in the progenitor strain lacking pAH88 (**Fig. 2F**). Upon the introduction of pAH88 to *S. coelicolor* M1152, ACT production was restored to the strain (**Fig. 2F**). It was found that in the three replicate populations at 1000, 2000 and 3000 generations that reintroduction of the ACT cluster identified pervasive negative epistasis across all strains, with no ACT production apparent except for one strain at the 1000 generation timepoint (**Fig. 2F**).

Examining genome sequence of the single ACT overproducing strain (Y1_P062_21; **Supp. Table S1**) indicates it possesses 25 mutations relative to the progenitor strain. This strain has substantial deletions at either end of the chromosome (Left-end of the chromosome 1-373,976 bp and right-end of the chromosome 7,773,260-8,494,261bp) resulting in the loss of 1019 genes. There are an additional four unique mutations in this strain. A missense mutation in SCO2160 (V500I; GTC-

>ATC) which encodes a putative large membrane protein, which to date has not been studied in *Streptomyces.* There is also a nonsense mutation in *cslA* (SCO2836) which encodes a cellulose synthase-like protein with a glycosyl transferase domain involved in hyphal tip growth and cell development [51]. Deletion of *clsA* is associated with reduced mycelial clumping and diffuse growth in liquid culture [50], which may also contribute to the fragmenting phenotype due to mutations in *matAB*. An intergenic mutation located between a small non-coding RNA (*scr5676*) [52] and *gabT* (SCO5676) encoding a 4-aminobutyrate aminotransferase resulting in the expansion of a G-tract (at position 6,080,415, G9->12). The positioning of the mutation downstream of these convergent genes appears unlikely to have any functional impact. There is a unique mutation in this strain in the pleiotropic regulator, *adpA* (W240*; discussed above), that results in the loss of the C-terminal helix-turn-helix DNA binding domain of AdpA [52]. Little is known about the role of the N-terminal domain of AdpA in antibiotic production and sporulation [53] or how this may function without a DNA binding domain. One mutation that is also present in this strain, and all strains sequenced in the Y1 replicate population, is an insertion of a C at 1119/1416 nucleotides resulting in a frameshift in *cmdB* (SC04127), a putative ATP/GTP binding protein that has previously been linked to negative regulation of ACT [54].

## Discussion

Experimental evolution is a powerful tool for exploring the mechanisms and dynamics of evolution [55]. In microbial experimental evolution the large populations coupled with rapid generation times allow the testing of evolutionary theories on long evolutionary time scales [14,55]. *Streptomyces* are excellent subjects for LTEE to study a range of adaptive behaviours given their remarkable hyphal growth habit, complex developmental lifecycle which are adaptations to growth in the terrestrial and ability to produce a plethora of bioactive molecules [20,56]. Growth in liquid culture results in relaxed selection on morphological and developmental traits given that sporulation in *S. coelicolor* does not occur in liquid culture and hyphae do not need to forage in a 3-dimensional environment for nutrients. These conditions are also representative of the fermentation conditions used in industry for the commercial production of antibiotics and other bioactive molecules.

Adaptive divergence in heterogeneous environments is well documented in bacteria [1,57,58]. Rainey and Travisano [1] demonstrated that a range of morphotypes emerge in *Pseudomonas* when grown in a heterogenous environment, but when cultures were mixed (homogenous environments) many of these morphotypes were absent. The emergence of developmental mutants during culture is well-known and frequent in *Streptomyces* reflecting the pleiotropic nature of many of the developmental genes identified to date, where genes associated with sporulation often also influence antibiotic production [59]. Prolonged liquid culture of *Streptomyces* in chemostats results in the emergence of *bld* mutants [60], although under these conditions, strains were not subjected to the dynamic fluctuations between nutrient sufficiency and starvation, which are key features of batch culture, which are known to trigger antibiotic biosynthesis [49,50] .

Reduction in the overall hyphal aggregate size likely relieves the nutrient limitation and mass transfer issues associated with growth of *Streptomyces* as cell aggregates [21,22]. Initially this led to increased fitness in the replicate populations, which appears to provide a trade off against hyphal lifestyle and development, with the parallel emergence of independent fragmenting hyphal and asporulent phenotypes. Whilst rapid growth is not always favoured in selective sweeps, the role of environmental fluctuation is thought to play a major role in the fixation of major phenotypic transitions [61]. The parallel emergence of fragmentation and loss of sporulation reflects phenotypes that are adaptative to the liquid environment when periodic selection for sporulation on solid medium in not encountered. The rapid, parallel emergence of the fragmentation phenotype would also suggest that this phenotype may favour rapid cell growth, and this is observed when individual clones are grown in the absence of competition. It is known that experimental lineages of *Escherichia coli* evolve to grow more rapidly over successive generations through increasing the efficiency of cellular metabolism [62–65]. The fragmenting phenotype of *Streptomyces* is associated with more rapid growth at the population level initially (1000 generations) along with the emergence of a range of morphotypes. Sequencing of strains identified pervasive mutations in loci known to affect fragmentation of hyphae in all strains, suggesting linkage to the phenotype. At the isolate level the divergence in fitness suggests that diversity is emerging within the cultures, even when strains are adapting to the same conditions [66]. If this is the case, then stochastic mutational accumulation within the replicate populations would likely lead to the appearance of metabolic diversity, and this was apparent in the three replicate lineages. Many of the mutations in primary metabolic processes link to transporters and regulators suggesting broad effects on metabolism. If random mutation and genetic drift was responsible for this, then carbon source utilisation would diversify in a random manner in the replicate populations. If patterns of carbon utilisation displayed parallel changes in replicate populations, it would suggest that antagonistic pleiotropy was occurring in the replicate populations [10]. These data suggest that both forces are likely at play under these conditions, where parallel increases in fitness can be seen for some carbon sources (Antagonistic pleiotropy) and non-parallel declines in carbon source utilisation can also be observed (mutation accumulation).

The *S*LTEE was designed to examine the role of epistasis in antibiotic production. This was facilitated using a strain that lacked four of the major antibiotic biosynthetic pathways [35], which allowed for the reintroduction of BGCs at specific points to understand how regulation changes under relaxed selection. Regulation of antibiotic biosynthesis in *Streptomyces* is multi-level, occurring at the global level and at the level of BGCs [49,50]. Upon reintroduction of the ACT BGC, it was found that negative epistasis was predominant, as has been observed previously for certain traits during adaptation [16,67,68]. This negative epistasis was parallel across genomes evolved in the absence of the ACT BGC. Genes of known function in antibiotic biosynthesis in the sequencing data mostly map to pleiotropic regulators, but there are several other genes present that are of unknown function suggesting there is plasticity in these pathways. These data also suggest that antagonistic interaction between global BGC regulation and within-BGC regulation of antibiotic biosynthesis is occurring. The discovery of this antagonistic interaction at the level of BGC regulation has not been documented previously in antibiotic biosynthesis.

This long-term evolution experiment in *Streptomyces* establishes a unique system to study how a hyphal, sporulating organism adapts to a novel environment. The experiment also facilitates the study of epistasis in antibiotic biosynthesis. The work demonstrates that adaptive radiation in *Streptomyces* results in the rapid emergence of multiple morphotypes within the population which exhibit hyphal fragmentation phenotypes as result of selection acting on hyphal aggregates, which limit nutrient uptake. Ecological opportunity maybe facilitating the divergence of strains with antagonistic pleiotropy evident from the parallelism observed in carbon utilisation between the replicate populations. The concomitant appearance of strains with developmental defects is due to relaxed selection on sporulation suggesting that accumulation of mutations occurs in genes required for development when sporulation is not permitted. Extensive negative epistasis was observed in the control of antibiotic biosynthesis when strains evolve without the ACT BGC, suggesting that antagonistic interactions may be at play between global and cluster-situated regulators. This on-going experimental system provides an excellent, multi-faceted model for the studying adaptation in *Streptomyces* and will generate important data that will find utility in ecology, evolution and has relevance to understanding industrial antibiotic production.

### CRediT (Contributor roles taxonomy)

Conceptualization – PAH.

Data curation - JTM, SV, LK.

Formal analysis - JTM, SV

PAH. Funding acquisition – PAH.

Investigation - JTM, SV, LK.

Methodology - JTM, SV, LK, PAH.

Project administration - JTM, PAH.

Supervision - JTM, PAH.

Validation - JTM, SV, PAH.

Visualization – JTM, SV

Writing – original draft – PAH.

Writing – review & editing - JTM, SV, LK, PAH.

All authors gave final approval for publication and agreed to be held accountable for the work performed therein.

### Ethics

This work did not require ethical approval from a human subject or animal welfare committee.

### Data accessibility

The data relevant to the conclusions of this paper and supplementary material is deposited online in Figshare (https://doi.org/10.6084/m9.figshare.29950388.v1).

### Declaration of AI use

We have not used AI-assisted technologies in creating this article.

## Acknowledgements

We would like to thank Dr Liam Rooney for help with microscopy and Dr Ryan Seipke for helpful discussions.

---

**Supplementary data 1** – All BactEXTRACT data output (growthParameters) for all Biolog PM1 data. Edited to include plate well annotations to specify carbon sources.

**Supplementary data 2** – Modified version **Supplementary data 1** where carbon source utilisation was filtered such that only those carbon sources have at least 1 replicate exhibiting classical bacterial growth profiles along with filtering columns for specific carbon sources.

**Supplementary figure 1 –** Additional examples of diversification of carbon source utilisation in the replicate populations over 3000 generations. Each point represents the growth of one of the replicate populations on a particular carbon source in the LTEE. Three carbon sources were chosen to illustrate different usage profiles as the population adapts to the medium through changes to specific growth rate, µ (h^-1^) Diversification of carbon source utilisation in the replicate populations over 3000 generations. Each point represents the growth of one of the replicate populations on a particular carbon source in the LTEE. Three carbon sources were chosen to illustrate different usage profiles as the population adapts to the medium through changes to specific growth rate, µ (h^-1^) 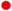 acetic acid; 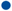 2-deoxyadenosine; 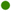 Tween 40.

**Supplementary table 1** – Breseq analysis of isolates indicating predicted mutations with addition of evidenced large deletions at start and end of chromosome. Highlighting classical developmental genes (green), Y1_P062_21 (act over producer) unique mutations (yellow) and multiple alleles/mutations in the same gene (red).

## References

1. Rainey PB, Travisano M. 1998 Adaptive radiation in a heterogeneous environment. Nature 394, 69–72. (doi:10.1038/27900)

2. Consuegra J, Plucain J, Gaffé J, Hindré T, Schneider D. 2017 Genetic Basis of Exploiting Ecological Opportunity During the Long-Term Diversification of a Bacterial Population. J. Mol. Evol. 85, 26–36. (doi:10.1007/s00239-017-9802-z)

3. Saxer G, Travisano M. 2016 Parallelism in adaptive radiations of experimental Escherichia coli populations. Evolution 70, 98–110. (doi:10.1111/evo.12841)

4. Saxer G, Doebeli M, Travisano M. 2010 The Repeatability of Adaptive Radiation During Long-Term Experimental Evolution of Escherichia coli in a Multiple Nutrient Environment. PLoS ONE 5, e14184. (doi:10.1371/journal.pone.0014184)

5. Kawecki TJ, Lenski RE, Ebert D, Hollis B, Olivieri I, Whitlock MC. 2012 Experimental evolution. Trends Ecol Evol 27, 547–560. (doi:10.1016/j.tree.2012.06.001)

6. Lenski RE. 2017 Experimental evolution and the dynamics of adaptation and genome evolution in microbial populations. ISME J. 11, 2181–2194. (doi:10.1038/ismej.2017.69)

7. Blount ZD, Lenski RE, Losos JB. 2018 Contingency and determinism in evolution: Replaying life’s tape. Science 362. (doi:10.1126/science.aam5979)

8. Jagdish T, Ba ANN. 2022 Microbial experimental evolution in a massively multiplexed and high-throughput era. Curr. Opin. Genet. Dev. 75, 101943. (doi:10.1016/j.gde.2022.101943)

9. Stroud JT, Ratcliff WC. 2025 Long-term studies provide unique insights into evolution. Nature 639, 589–601. (doi:10.1038/s41586-025-08597-9)

10. Cooper VS, Lenski RE. 2000 The population genetics of ecological specialization in evolving Escherichia coli populations. Nature 407, 736–739. (doi:10.1038/35037572)

11. Cooper VS. 2014 The Origins of Specialization: Insights from Bacteria Held 25 Years in Captivity. PLoS Biol. 12, e1001790. (doi:10.1371/journal.pbio.1001790)

12. Maharjan R, McKenzie C, Yeung A, Ferenci T. 2013 The basis of antagonistic pleiotropy in hfq mutations that have opposite effects on fitness at slow and fast growth rates. Heredity 110, 10–18. (doi:10.1038/hdy.2012.46)

13. Ruelens P, Wynands T, Visser JAGM de. 2023 Interaction between mutation type and gene pleiotropy drives parallel evolution in the laboratory. Philos. Trans. R. Soc. B 378, 20220051. (doi:10.1098/rstb.2022.0051)

14. Lenski RE. 2017 Experimental evolution and the dynamics of adaptation and genome evolution in microbial populations. Isme J 11, 2181–2194. (doi:10.1038/ismej.2017.69)

15. Lenski RE. 2017 What is adaptation by natural selection? Perspectives of an experimental microbiologist. Plos Genet 13, e1006668. (doi:10.1371/journal.pgen.1006668)

16. Khan AI, Dinh DM, Schneider D, Lenski RE, Cooper TF. 2011 Negative Epistasis Between Beneficial Mutations in an Evolving Bacterial Population. Science 332, 1193–1196. (doi:10.1126/science.1203801)

17. Kryazhimskiy S, Rice DP, Jerison ER, Desai MM. 2014 Global epistasis makes adaptation predictable despite sequence-level stochasticity. Science 344, 1519–1522. (doi:10.1126/science.1250939)

18. Flärdh K, Buttner MJ. 2009 Streptomyces morphogenetics: dissecting differentiation in a filamentous bacterium. Nature Reviews Microbiology 7. (doi:10.1038/nrmicro1968)

19. Elliot MA, Buttner MJ, Nodwell JR. 2008 Multicellular Development in Streptomyces. In (ed DE Whitworth), pp. 419–439. ASM.

20. Hoskisson PA, Barona-Gómez F, Rozen DE. 2024 Phenotypic heterogeneity in Streptomyces colonies. Curr. Opin. Microbiol. 78, 102448. (doi:10.1016/j.mib.2024.102448)

21. Nieminen L, Webb S, Smith MCM, Hoskisson PA. 2013 A Flexible Mathematical Model Platform for Studying Branching Networks: Experimentally Validated Using the Model Actinomycete, Streptomyces coelicolor. PLoS ONE 8. (doi:10.1371/journal.pone.0054316)

22. Dissel D van, Claessen D, Roth M, Wezel GP van. 2015 A novel locus for mycelial aggregation forms a gateway to improved Streptomyces cell factories. Microb Cell Fact 14, 44. (doi:10.1186/s12934-015-0224-6)

23. Bentley SD et al. 2002 Complete genome sequence of the model actinomycete Streptomyces coelicolor A3(2). Nature 417, 141–7. (doi:10.1038/417141a)

24. Schniete JK, Cruz-Morales P, Selem-Mojica N, Fernández-Martínez LT, Hunter IS, Barona- Gómez F, Hoskisson PA. 2018 Expanding Primary Metabolism Helps Generate the Metabolic Robustness To Facilitate Antibiotic Biosynthesis in Streptomyces. Mbio 9, e02283–17. (doi:10.1128/mbio.02283-17)

25. Lioy VS et al. 2021 Dynamics of the compartmentalized Streptomyces chromosome during metabolic differentiation. Nat Commun 12, 5221. (doi:10.1038/s41467-021-25462-1)

26. Gravius B, Bezmalinović T, Hranueli D, Cullum J. 1993 Genetic instability and strain degeneration in Streptomyces rimosus. Appl Environ Microb 59, 2220–2228. (doi:10.1128/aem.59.7.2220-2228.1993)

27. Altenbuchner J, Cullum J. 1984 DNA amplification and an unstable arginine gene in Streptomyces lividans 66. Mol. Gen. Genet. MGG 195, 134–138. (doi:10.1007/bf00332735)

28. Leblond P, Demuyter P, Moutier L, Laakel M, Decaris B, Simonet JM. 1989 Hypervariability, a new phenomenon of genetic instability, related to DNA amplification in Streptomyces ambofaciens. J. Bacteriol. 171, 419–423. (doi:10.1128/jb.171.1.419-423.1989)

29. Fischer G, Wenner T, Decaris B, Leblond P. 1998 Chromosomal arm replacement generates a high level of intraspecific polymorphism in the terminal inverted repeats of the linear chromosomal DNA of Streptomyces ambofaciens. Proc. Natl. Acad. Sci. 95, 14296–14301. (doi:10.1073/pnas.95.24.14296)

30. Hoff G, Bertrand C, Piotrowski E, Thibessard A, Leblond P. 2018 Genome plasticity is governed by double strand break DNA repair in Streptomyces. Sci. Rep. 8, 5272. (doi:10.1038/s41598-018-23622-w)

31. Bury-Moné S, Thibessard A, Lioy VS, Leblond P. 2023 Dynamics of the Streptomyces chromosome: chance and necessity. Trends Genet. (doi:10.1016/j.tig.2023.07.008)

32. Zhang Z, Shitut S, Claushuis B, Claessen D, Rozen DE. 2022 Mutational meltdown of putative microbial altruists in Streptomyces coelicolor colonies. Nat Commun 13, 2266. (doi:10.1038/s41467-022-29924-y)

33. Zhang Z, Du C, Barsy F de, Liem M, Liakopoulos A, Wezel GP van, Choi YH, Claessen D, Rozen DE. 2020 Antibiotic production in Streptomyces is organized by a division of labor through terminal genomic differentiation. Sci. Adv. 6, eaay5781. (doi:10.1126/sciadv.aay5781)

34. Traxler MF, Rozen DE. 2022 Ecological drivers of division of labour in Streptomyces. Curr Opin Microbiol 67, 102148. (doi:10.1016/j.mib.2022.102148)

35. Gomez-Escribano JP, Bibb MJ. 2011 Engineering Streptomyces coelicolor for heterologous expression of secondary metabolite gene clusters. Microb Biotechnol 4, 207–215. (doi:10.1111/j.1751-7915.2010.00219.x)

36. Feeney MA et al. 2022 ActinoBase: tools and protocols for researchers working on Streptomyces and other filamentous actinobacteria. Microb Genom 8. (doi:10.1099/mgen.0.000824)

37. Schindelin J et al. 2012 Fiji: an open-source platform for biological-image analysis. Nat. Methods 9, 676–682. (doi:10.1038/nmeth.2019)

38. Dénéréaz J, Veening J-W. 2024 BactEXTRACT: an R Shiny app to quickly extract, plot and analyse bacterial growth and gene expression data. Access Microbiol. 6, 000742.v3. (doi:10.1099/acmi.0.000742.v3)

39. Larcombe DE, Braes RE, Croxford JT, Wilson JW, Figurski DH, Hoskisson PA. 2024 Sequence and origin of the Streptomyces intergenetic-conjugation helper plasmid pUZ8002. Access Microbiol. 6. (doi:10.1099/acmi.0.000808.v3)

40. Deatherage DE, Barrick JE. 2014 Engineering and Analyzing Multicellular Systems, Methods and Protocols. Methods Mol. Biol. 1151, 165–188. (doi:10.1007/978-1-4939-0554-6_12)

41. Schlimpert S, Elliot MA. 2023 The Best of Both Worlds—Streptomyces coelicolor and Streptomyces venezuelae as Model Species for Studying Antibiotic Production and Bacterial Multicellular Development. J. Bacteriol. 205, e00153–23. (doi:10.1128/jb.00153-23)

42. Buckling A, Wills MA, Colegrave N. 2003 Adaptation Limits Diversification of Experimental Bacterial Populations. Science 302, 2107–2109. (doi:10.1126/science.1088848)

43. Poltak SR, Cooper VS. 2011 Ecological succession in long-term experimentally evolved biofilms produces synergistic communities. ISME J. 5, 369–378. (doi:10.1038/ismej.2010.136)

44. Kovács ÁT. 2023 Colony morphotype diversification as a signature of bacterial evolution. microLife 4, uqad041. (doi:10.1093/femsml/uqad041)

45. Takemoto N, Numata I, Su’etsugu M, Miyoshi-Akiyama T. 2018 Bacterial EndoMS/NucS acts as a clamp-mediated mismatch endonuclease to prevent asymmetric accumulation of replication errors. Nucleic Acids Res. 46, 6152–6165. (doi:10.1093/nar/gky481)

46. Dagva O, Thibessard A, Lorenzi J-N, Labat V, Piotrowski E, Rouhier N, Myllykallio H, Leblond P, Bertrand C. 2024 Correction of non-random mutational biases along a linear bacterial chromosome by the mismatch repair endonuclease NucS. Nucleic Acids Res. 52, 5033–5047. (doi:10.1093/nar/gkae132)

47. Vasi F, Travisano M, Lenski RE. 1994 Long-Term Experimental Evolution in Escherichia coli. II. Changes in Life-History Traits During Adaptation to a Seasonal Environment. Am. Nat. 144, 432–456. (doi:10.1086/285685)

48. Lenski RE, Rose MR, Simpson SC, Tadler SC. 1991 Long-Term Experimental Evolution in Escherichia coli. I. Adaptation and Divergence During 2,000 Generations. Am. Nat. 138, 1315– 1341. (doi:10.1086/285289)

49. Bibb MJ. 2005 Regulation of secondary metabolism in streptomycetes. Current Opinion in Microbiology 8. (doi:10.1016/j.mib.2005.02.016)

50. Hoskisson PA, Fernández-Martínez LT. 2018 Regulation of specialised metabolites in Actinobacteria – expanding the paradigms. Env Microbiol Rep 10, 231–238. (doi:10.1111/1758-2229.12629)

51. Xu H, Chater KF, Deng Z, Tao M. 2008 A Cellulose Synthase-Like Protein Involved in Hyphal Tip Growth and Morphological Differentiation in Streptomyces. J. Bacteriol. 190, 4971–4978. (doi:10.1128/jb.01849-07)

52. Swiercz JP, Hindra, Bobek J, Bobek J, Haiser HJ, Berardo CD, Tjaden B, Elliot MA. 2008 Small non-coding RNAs in Streptomyces coelicolor. Nucleic Acids Res. 36, 7240–7251. (doi:10.1093/nar/gkn898)

53. Yamazaki H, Tomono A, Ohnishi Y, Horinouchi S. 2004 DNA-binding specificity of AdpA, a transcriptional activator in the A-factor regulatory cascade in Streptomyces griseus. Molecular Microbiology 53. (doi:10.1111/j.1365-2958.2004.04153.x)

54. Xu Z, Li Y, Wang Y, Deng Z, Tao M. 2019 Genome-Wide Mutagenesis Links Multiple Metabolic Pathways with Actinorhodin Production in Streptomyces coelicolor. Appl. Environ. Microbiol. 85, e03005–18. (doi:10.1128/aem.03005-18)

55. McDonald MJ. 2019 Microbial Experimental Evolution – a proving ground for evolutionary theory and a tool for discovery. EMBO Rep. 20, EMBR201846992. (doi:10.15252/embr.201846992)

56. Claessen D, Jong W de, Dijkhuizen L, Wösten H. 2006 Regulation of Streptomyces development: reach for the sky! Trends in Microbiology 14. (doi:10.1016/j.tim.2006.05.008)

57. Herron MD, Doebeli M. 2013 Parallel Evolutionary Dynamics of Adaptive Diversification in Escherichia coli. Plos Biol 11, e1001490. (doi:10.1371/journal.pbio.1001490)

58. Houte S van, Padfield D, Gómez P, Luján AM, Brockhurst MA, Paterson S, Buckling A. 2021 Compost spatial heterogeneity promotes evolutionary diversification of a bacterium. J. Evol. Biol. 34, 246–255. (doi:10.1111/jeb.13722)

59. Chandra G, Chater KF. 2014 Developmental biology of Streptomyces from the perspective of 100 actinobacterial genome sequences. FEMS Microbiology Reviews 38, 345–379. (doi:10.1111/1574-6976.12047)

60. Butler PR, Brown M, Oliver SG. 1996 Improvement of antibiotic titers from Streptomyces bacteria by interactive continuous selection. Biotechnol. Bioeng. 49, 185–196. (doi:10.1002/(sici)1097-0290(19960120)49:2<185::aid-bit7>3.0.co;2-m)

61. Zhu M, Dai X. 2024 Shaping of microbial phenotypes by trade-offs. Nat. Commun. 15, 4238. (doi:10.1038/s41467-024-48591-9)

62. Marshall DJ, Malerba M, Lines T, Sezmis AL, Hasan CM, Lenski RE, McDonald MJ. 2022 Long-term experimental evolution decouples size and production costs in Escherichia coli. Proc. Natl. Acad. Sci. 119, e2200713119. (doi:10.1073/pnas.2200713119)

63. Lenski RE, Travisano M. 1994 Dynamics of adaptation and diversification: a 10,000-generation experiment with bacterial populations. Proc. Natl. Acad. Sci. 91, 6808–6814. (doi:10.1073/pnas.91.15.6808)

64. Grant NA, Magid AA, Franklin J, Dufour Y, Lenski RE. 2021 Changes in Cell Size and Shape during 50,000 Generations of Experimental Evolution with Escherichia coli. J Bacteriol 203. (doi:10.1128/jb.00469-20)

65. Lenski RE et al. 2015 Sustained fitness gains and variability in fitness trajectories in the long- term evolution experiment with Escherichia coli. Proc. R. Soc. B: Biol. Sci. 282, 20152292. (doi:10.1098/rspb.2015.2292)

66. Maddamsetti R, Grant NA. 2020 Divergent Evolution of Mutation Rates and Biases in the Long- Term Evolution Experiment with Escherichia coli. Genome Biol. Evol. 12, 1591–1603. (doi:10.1093/gbe/evaa178)

67. Chou H-H, Chiu H-C, Delaney NF, Segrè D, Marx CJ. 2011 Diminishing Returns Epistasis Among Beneficial Mutations Decelerates Adaptation. Science 332, 1190–1192. (doi:10.1126/science.1203799)

68. Zee PC, Velicer GJ. 2017 Parallel emergence of negative epistasis across experimental lineages. Evolution 71, 1088–1095. (doi:10.1111/evo.13190)

